# Insights into the binding mode of MEK type-III inhibitors. A step towards discovering and designing allosteric kinase inhibitors across the human kinome

**DOI:** 10.1101/076711

**Authors:** Zheng Zhao, Lei Xie, Philip E. Bourne

**Affiliations:** National Center for Biotechnology Information, National Library of Medicine, National Institute of Health, Bethesda, MD 20892, USA; Department of Computer Science, Hunter College, The City University of New York, NY 10065, USA; The Graduate Center, The City University of New York, NY 10016, USA; Office of the Director, National Institutes of Health, Bethesda, MD 20892, USA

## Abstract

Protein kinases are critical drug targets for treating a large variety of human diseases. Type-I and type-II kinase inhibitors frequently exhibit off-target toxicity or lead to mutation acquired resistance. Two highly specific allosteric type-III MEK-targeted drugs, Trametinib and Cobimetinib, offer a new approach. Thus, understanding the binding mechanism of existing type-III kinase inhibitors will provide insights for designing new type-III kinase inhibitors. In this work we have systematically studied the binding mode of MEK-targeted type-III inhibitors using structural systems pharmacology and molecular dynamics simulation. Our studies provide detailed sequence, structure, interaction-fingerprint, pharmacophore and binding-site information on the binding characteristics of MEK type-III kinase inhibitors. We propose that the helix-folding activation loop is a hallmark allosteric binding site for type-III inhibitors. Subsequently we screened and predicted allosteric binding sites across the human kinome, suggesting other kinases as potential targets suitable for type-III inhibitors. Our findings will provide new insights into the design of potent and selective kinases inhibitors.

**Author Summary:** Human protein kinases represent a large protein family relevant to many diseases, especially cancers, and have become important drug targets. However, developing the desired selective kinase-targeted inhibitors remain challenging. Kinase allosteric inhibitors provide that opportunity, but, to date, few have been designed other than MEK inhibitors. To more efficiently develop kinase allosteric inhibitors, we systematically studied the binding mode of the MEK type-III allosteric kinase inhibitors using structural system pharmacology and molecular dynamics approaches. New insights into the binding mode and mechanism of type-III inhibitors were revealed that may facilitate the design of new prospective type-III kinase inhibitors.

## Introduction

Kinases are phosphorylation enzymes that catalyze the transfer of phosphate groups from ATP to specific substrates and are critical in most cellular life processes [1, 2]. Abnormal kinase regulation, which leads to signal disruption and cell deregulation, is implicated in many diseases, particularly cancers [3]. Thus a number of kinase-targeted small molecule inhibitors have been developed and are important in anti-cancer therapy [4]. Through July 2016, 30 small molecule kinase inhibitors [5, 6] have been approved by the US Food and Drug Administration (FDA) for the treatment of cancers or other diseases (http://www.fda.gov/), and additional inhibitors are undergoing clinical trials [7, 8].

However, reported off-target toxicities and acquired-mutation resistance [9] require kinase-targeted inhibitors of lower dose and higher specificity. Three types of targeted kinase inhibitors, type-I, type-II and type-III, have been developed [10, 11]. Type-I inhibitors are ATP-competitive and occupy the ATP-binding binding pocket, a highly conserved kinase catalytic scaffold with strong binding affinity for ATP. The difficulty of achieving selectivity combined with mutation-induced resistance has led to the development of type-II and type-III inhibitors [12]. Type-II inhibitors bind to the inactive kinase and not only occupy the ATP-binding pocket, but extend to occupy the adjacent less-conserved allosteric site across the DFG segment, which is accessible in the DFG-out inactive kinase conformation [13]. Most type-I and type-II inhibitors bind to multiple targets[9]. Type-III inhibitors are unique in that they occupy highly specific allosteric sites as exemplified by the type-III MEK inhibitors [14, 15]. Besides the type-III MEK inhibitors, a few type-III inhibitors are reported[16] including the BCR-ABL inhibitors GNF2 and ABL001 [17], the pan-AKT inhibitor MK-2206 [18] and the mutant-selective EGFR allosteric inhibitor EAI045 [19]. The reported evidence so far suggests that type-III inhibitors are highly selective and effective even if the kinase or signaling pathway has undergone drug-resistance residue mutation [16, 19]. For example, an FDA-approved type-III MEK kinase inhibitor, Cobimetinib (IC50:0.9 nM), can overcome the resistance from the typical BRAF V600E mutation seen in melanoma [15]. However, thus far, there is no systematic means for identifying the preferred characteristics for specific type-III inhibitors [8]. Existing type-III kinase inhibitors mainly target MEK[16]. Thus it is important to understand the molecular characteristics of the interaction between MEK kinase and type-III kinase inhibitors, so as to extend the design of type-III kinase inhibitors and develop the type-III targeting libraries.

In this work we have integrated our structural systems biology strategy and molecular dynamics simulation methods to gain insights into type-III kinase inhibitors and their binding modes with kinases across the human kinome. Our structural system biology strategy harnesses different omics data to compare and discover the gene-level, protein-level and structure-level information on protein-ligand interactions [20]. We have successfully applied this strategy to drug design and discovery for different target-drug kinase systems and the Ebola virus proteome [5, 21]. With increased computing power and more efficient algorithms, molecular dynamics (MD) simulation is now becoming a routine tool for drug design, accounting for the reality of a flexible target structure and flexible target-drug binding [22]. In this paper we performed detailed MEK kinase-drug function-site interactional fingerprint analysis using our structural system biology strategy. We also performed two MD simulations up to 1.2 μs in an explicit water box to obtain insights into the behavior of flexible MEK kinases as targets, with and without a representative ligand, Cobimetinib [23]. By comparing the structural trajectories between MEK with and without ligand, we determined the structural flexibility and interactional network for type-III inhibitor binding to MEK kinases. In addition, we delved into the details of genetic structural, pharmacophore and mechanistic understanding of MEK kinase-drug binding. Finally, we explored the whole human kinome to identify potentially new opportunities for type-III inhibitor drug design.

## Results

### Binding modes of crystallized ligand-bound MEK complexes

We obtained the binding characteristics of ligand-bound MEK complexes as shown in Fig 1. Fig 1a illustrates the alignment of the 28 catalytic kinase domains of MEK bound to the type III inhibitors shown in the same allosteric binding site. We calculated the detailed interactions between MEK and the ligand using the Fs-IFP method (Fig 1b). The highly conserved interactions between the respective ligands and MEK include K97, L115, L118, V127, M143, C^207^DFGVS^212^, I215 and M219 (Fig 1b, and 1c). These conserved interactions can be divided into three spatial regions.

**Fig 1.**
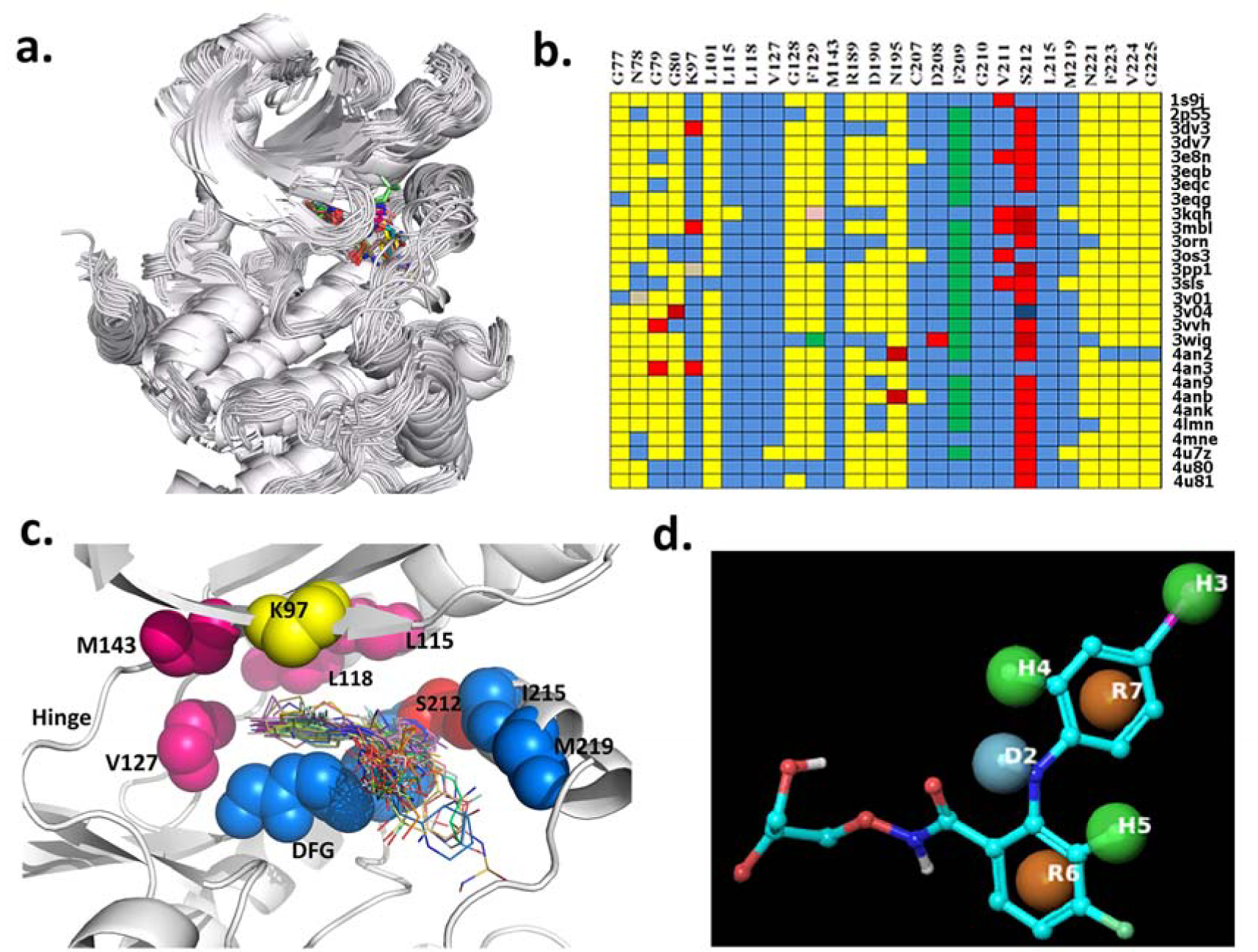
Binding characters of MEK-ligand complexes. a) All MEK-ligand complex structures aligned using SMAP. b) Encoding all MEK-ligand interactions. Every row represents one MEK-ligand interaction of one complex structure, and every column represents the interactions between the same amino acid and the bound ligand in different complex structures. Different colors represent the different types of fingerprint interactions: yellow, no interaction; blue, apolar interaction; red, apolar interaction and hydrogen bond interaction (protein as donor); deep red, hydrogen bond interaction (protein as donor); pink, polar interaction and aromatic interaction; and grey, apolar interaction and hydrogen bond interaction (protein as acceptor). c) Spatial representation of MEK-ligand interactions. d) Pharmacophore modeling: H, hydrophobic group; R, aromatic ring; D, hydrogen-bond donor.

The first region is the hydrophobic sub-pocket consisting of L115, L118, V127 and M143 as shown in purple in Fig 1c. All interactions are apolar (Fig 1b). Correspondingly, all ligands have hydrophobic groups that can be accommodated in the sub-pocket. For example, Cobimetinib has a 2-fluoro-4-iodoanilino fragment, as shown in purple in Fig 2 (4an2), which is a well-known hydrophobic pocket binder and accommodates the hydrophobic sub-pocket with hydrophobic contacts as shown in Fig 1c. Other ligands also have the same or similar fragments (Fig 2, the fragment in purple) so as to achieve the high binding affinity of the conserved sub-pocket. Pharmacophore modeling, as shown in Fig 1d, also revealed similar patterns with three common hydrophobic groups, H3, H4 and R7.

**Fig 2.**
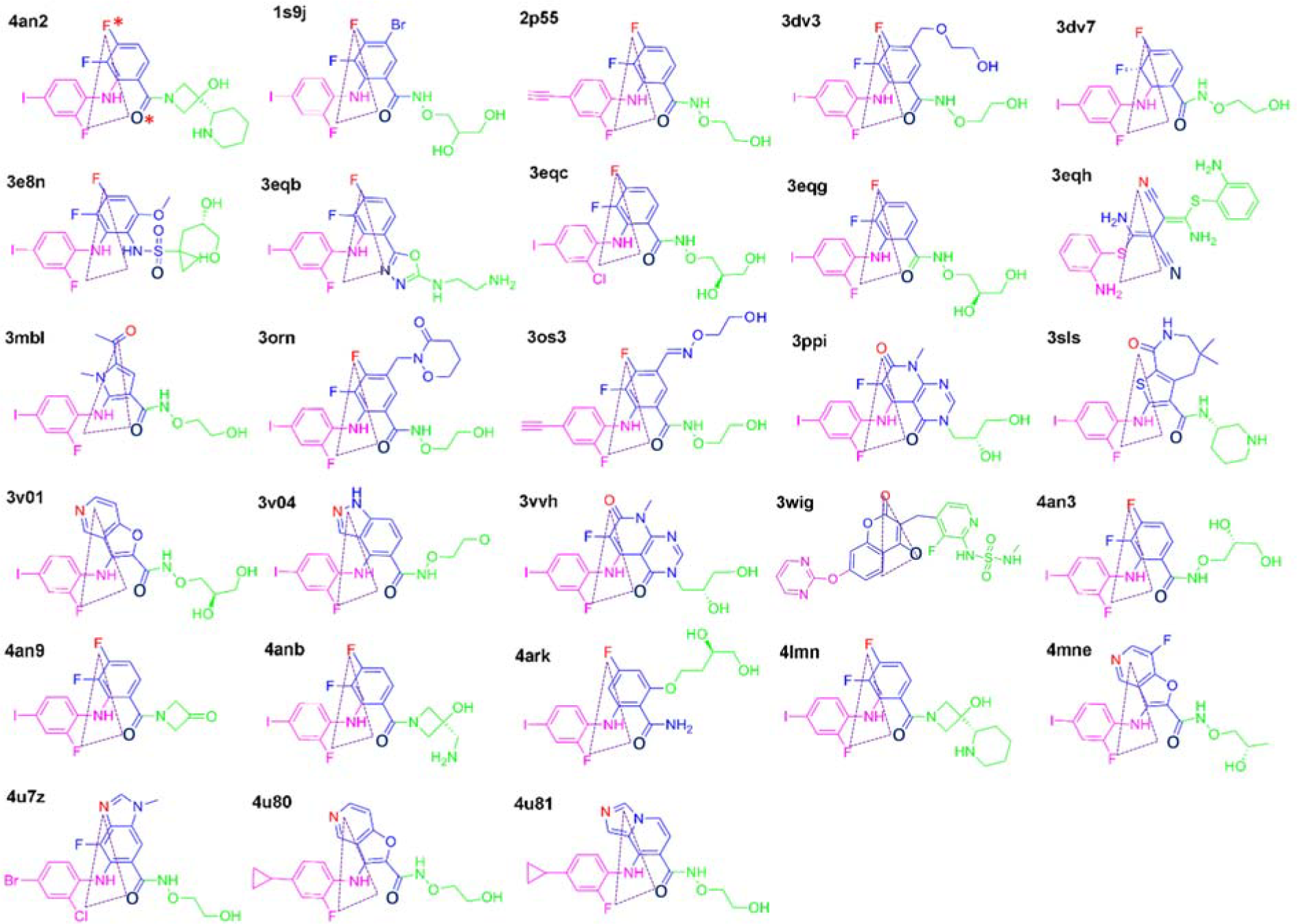
Ligands from the 28 MEK-ligand complex structures. The triangles highlight the conserved atoms that characterize these MEK Type-III inhibitors in 3D space.

The second region is K97, an important catalytic residue, located at the roof of the binding pocket (Fig 1c, yellow color). K97 has a conserved interaction with the oxygen atom O* (marked in Fig 2, dark blue color) in the ligands. Moreover, the molecular moiety O* is conserved, shown in the same position for other compounds in dark blue, either as an oxygen or nitrogen atom (Fig 2, 3eqb and 3eqh). Pharmacophore modeling (Fig 1d) is consistent with one donor-type hydrogen bond, D2.

The third region consisting of C^207^DFGVS^212^, I215 and M219 forms a loop and a helix that acts like an arm to accommodate the inhibitor to the kinase active site. DFG is directly involved in kinase catalytic activity across the family. I215 and M219 are located at the activation loop. Interestingly in MEK the activation loop is a conserved helix which forms the allosteric site. In most MEK structures, S212 forms a conserved interaction (Fig 1b, S212 column, red color) including an apolar and a hydrogen-bond interaction. Only in the structures 1s9j, 3egg, 3oss, 4v04 and 4an3, does S212 provide only a polar interactions to the corresponding ligand. This conserved interaction is consistent with experiment which shows S212 plays a key role in phosphorylation by RAF [15]. In this third region all active ligands have one atom, F, N or O that interacts with the backbone of S212 (shown as F* in red, Fig 2, 4an2). Pharmacophore modeling (Fig 1d, H5 and R6) illustrates that all ligands have common features in their interaction with S212.

Taken together, the aforementioned three regions make major contributions to ligand binding in the allosteric pocket. Summarizing Fig 1d, the hydrophobic heads (H3, H4 and R7) accommodate the hydrophobic sub-pocket and D2 and H5 interact with the roof amino acid K97 and S212 of the loop, respectively. It is expected that an active MEK inhibitor would have these chemical functional groups or similar. Furthermore, in 3D space, these atoms are spatially conserved, as shown by the triangles in Fig 2 for D2, H4 and H5, suggesting that the design of MEK allosteric inhibitors should follow this pharmacophore. Similar conserved spatial requirements have been reported in the design of other allosteric inhibitors [19]. Of course, besides the conserved pharmacophores, different inhibitors may use specific interactions involving other amino acids to achieve selectivity, as shown in Fig 1b. Specifically, the solvent exposed part of the inhibitor, shown in green in Fig 2, can be changed using different chemical group or atoms. The pharmacophore does not reveal common features for the solvent exposed part (Fig 1d). However, the difference in the solvent exposed part of the inhibitor also shows the different inhibition ability for MEK, such as shown the structure-activity relationship in the process of optimizing the compounds 2-19 toward Cobimetinib [14]. Interestingly, the solvent exposed parts of the inhibitors are right next to the γ-phosphate of ATP. The co-crystal MEK structure with ACP, which is an ATP analogue, and Cobimetinib (pdb id: 4an2) was shown in Fig S1, in which the γ-phosphate of the ACP interacts with the solvent exposed parts of Cobimetinib. It is clear that the interactions disturb the functional conformation of the γ-phosphate of ATP and the substrate being hosphorylated, which results in the redistribution of electron density in the active site, thereby reducing MEK enzymatic activity.

### MEK structural flexibility and insights into the binding mechanism

Beyond the static PDB structures we considered how type-III inhibitors induce MEK structural flexibility. We performed two 0.6 μs MD simulations for MEK kinase with and without the inhibitor bound, respectively. The Cα-atom RMSD is shown in Fig S2. The Cα-RMSDs of the apo and holo structures are similar. However, their Cα-atom fluctuations show significant difference flexibility, especially in the ligand-bound regions as shown in Fig 3a, where we compared MEK structural flexibility before and after binding using the two last 0.5 μs equilibrated MD trajectories. For apo MEK the flexibility change mainly comes from the P-loop, the activation loop and the C-terminal lobe as shown in Fig 3a. The corresponding collective motion as inferred from the first principal component of PCA is shown in Fig 3b. As a comparison, in the MEK-Cobimetinib complex, the main fluctuations come from the parts of the C-helix, the sequences following the activation loop and the C-terminal lobe (Fig 3a). The obvious difference before and after binding inhibitor is that the collective motions of the P-loop and activation loop have undergone a substantial reduction in the MEK-Cobimetinib complex (Fig 3c). However, the flexibility of the C-helix has significantly increased in apo MEK. Similar to other kinases, the P-loop contributes to conformational flexibility and plays an important role in binding and recognizing phosphoryl moieties [24]. Moreover, this flexible P-loop motif, along with other beta-sheets and helixes, generally form a pocket into which the phosphate groups can insert [25, 26]. Like the P-loop, the activation loop shows a similar change in flexibility, before and after Cobimetinib binding. Thus, MEK is induced to fit the inhibitor and the resulting increased rigidity in the activation-loop reduces MEK activity. Within the kinases the C-helix, Lys97 and DFG peptide contribute to form the ATP binding site needed for enzyme catalysis, including a hydrogen bond between the C-helix and Lys97. Upon Cobimetinib binding MEK enters an inactive state and the hydrogen bond is broken. Subsequently, the C-helix is more flexible than in the active state [8, 27]. In addition, the activation loop is a helix in MEK. However, in most kinases, the activation loop is a flexible loop. This implies that the helix is responsible for forming the binding allosteric pocket that fits the type-III inhibitor. Thus consideration of the nature of the adopted helix as part of the activation loop is an important consideration when designing type-III inhibitors.

**Fig 3.**
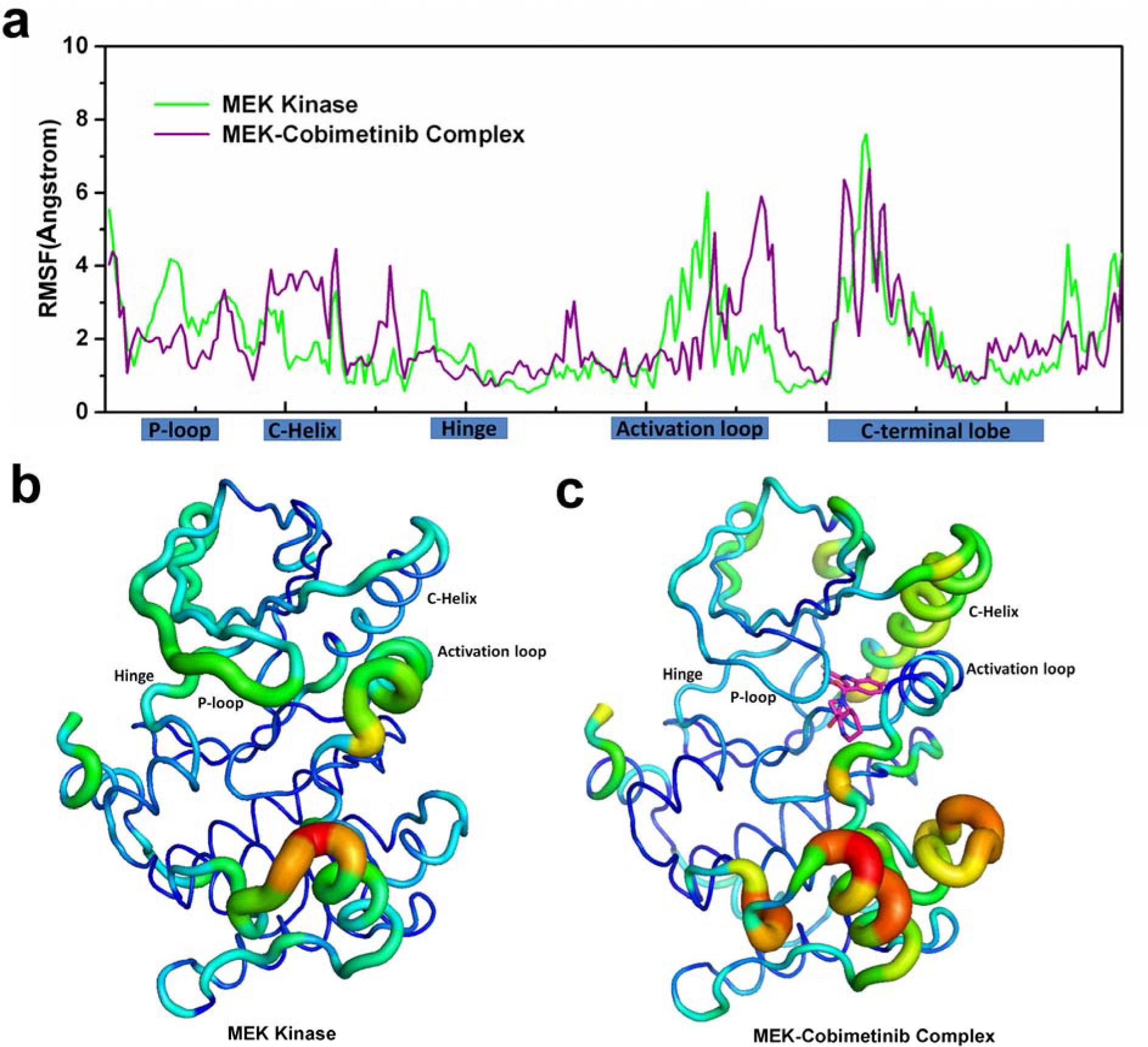
RMSF profiles and PCA projection. a) The RMSF profiles from the last 0.5 μm equilibrated MD trajectories of MEK and the MEK-Cobimetinib complex. Some secondary structure elements are shown on the abscissa. b,c) The Ca-atom projection along the first principal component. The displacements are shown as color-coded tubes from blue (small displacement) to orange (large displacement) for MEK and the MEK-Cobimetinib complex, respectively.

### Conserved interactions with inhibitors from S212 and K97

As aforementioned, the inhibitors derived from PDB MEK structures have a similar core and common functional groups forming a conserved spatial triangular arrangement (Fig 2). Correspondingly, in the MEK-Cobimetinib MD trajectories the conserved interactions between MEK and respective inhibitors were also evaluated. Two key interactions between S212 and K97 and the inhibitors were highlighted (Fig 4). The interaction between the backbone nitrogen atom of S212 and the F* atom of the inhibitor is shown in red; from the probability distribution (Fig 4b) the center of the peak is at 3.1 Å, which suggests that a hydrogen bond interaction is conserved at all times to maintain the binding affinity and restrain the flexible movement of S212 thereby hindering MEK phosphorylation by RAF. This observed hydrogen bond interaction is in agreement with the reported experimental results [15]. The O* atom is another conserved polar atom contributing to the effective binding. As shown in Fig 4, green color, the position of the peak in the probability distribution is at approximately 3.0 Å, which agrees with the distance found in PDB crystal structures such as 4lmn [14, 15]. This distance suggests that there is a strong hydrogen-bond interaction between the atom O* of the ligand and the ε-amino group of the lysine (K97), which contributes to forming the catalytic center [28]. Thus, the hydrogen-bond interaction blocks the salt-bridge interaction between the ε-amino group of the lysine and the E114 of the C-Helix. Importantly, blocking the salt-bridge interaction results in an inactive state [29] and increased flexibility of the C-Helix. The O* atom of the inhibitor, the ε-amino group of the lysine K97 and the carboxyl group of the aspartic acid of DFG form the pseudo catalytic center and deactivate the kinase catalytic function in the MEK/ERK pathway [30].

**Fig 4.**
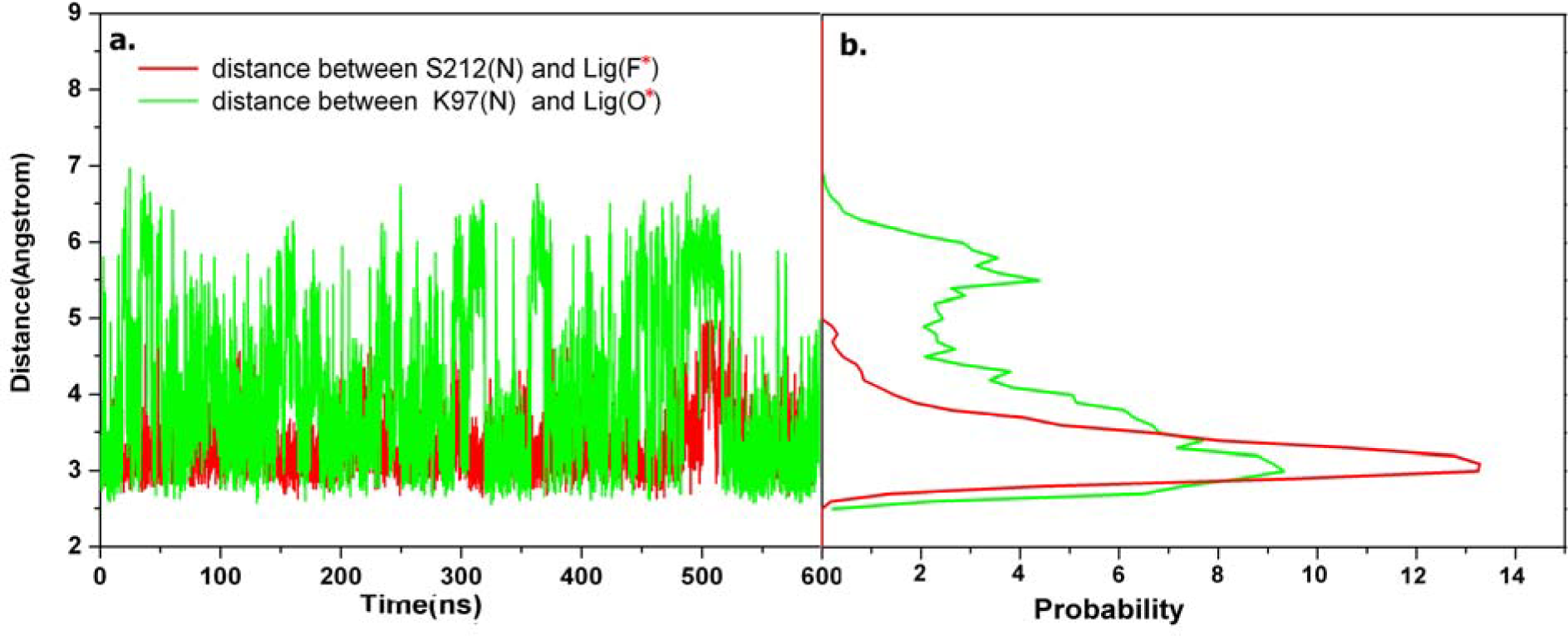
Conserved interatomic interactions between MEK and the ligand. a) Interatomic distance for every conformation in the MD trajectory; b) The probability distribution of interatomic distances.

### Ranking similar binding sites to MEK using SMAP across the structural kinome

To design type-III inhibitors for any given kinase target, establishing similar binding pockets to MEK is an important step. Using SMAP [31–33], we search for similar binding sites across the human kinome. Using the threshold of more than 55% similarity, three crystal structures with MEK-similar binding pockets were found (pdb ids 2yix, 4pp7 and 4wo5). Herein, one structure belongs to the kinase P38a [34] and two others belong to the kinase BRAF [2]. All three structures contain a helix-folding activation loop. We aligned the sequences of MEK, P38a and BRAF with particular attention to the activation loop. The similarity around them is not high as shown in Fig S3, where the activation loops are marked with a rectangle. These results imply that kinases with similar secondary structure in their activation loops have the potential to be inhibited by type-III inhibitors even though their global sequences do not have high similarity with MEK. Stated another way, a helix in the activation loop provides structural insight into designing type-III inhibitors and caused us to investigate which kinases can potentially form a helix in the activation loop.

### Predicting helix-folding activation loops across the human kinome

Developing type-III inhibitors needs to be validated not only by biological activity assay but also through analysis of an appropriate 3D complex structure [8]. Unfortunately, the type-III binding pocket is often not present in the 3D complex structure when there is no specific type-III inhibitor co-crystallized [8]. To obtain the MEK-similar allosteric site in other kinases, we predicted the MEK-like secondary structure of the activation loop that may critical for the binding pose of the type-III inhibitor [35]. Based on amino acid sequences of all kinases, we determined the top 15 kinases that have a potential allosteric site suitable for type-III inhibitors [8] as shown in Fig 5 and detailed in Table 1. The top 15 kinases are mostly located within the STE group forming the MAPK cascade. Interesting, at the level of sequence, their activation loops don't have high similarity to MEK. However, at the level of second structure, like MEK, the corresponding activation loops of the 15 kinases may be folded as a helix, as validated by a newly released X-ray structure of the kinase MAP2K4 (pdb id 3alo) [36], in which the crystallized activation loop shows a helix folding mode. Thus the helix folding of activation loops potentially acts like an anchor to accommodate the type-III inhibitor in the active site.

**Fig 5.**
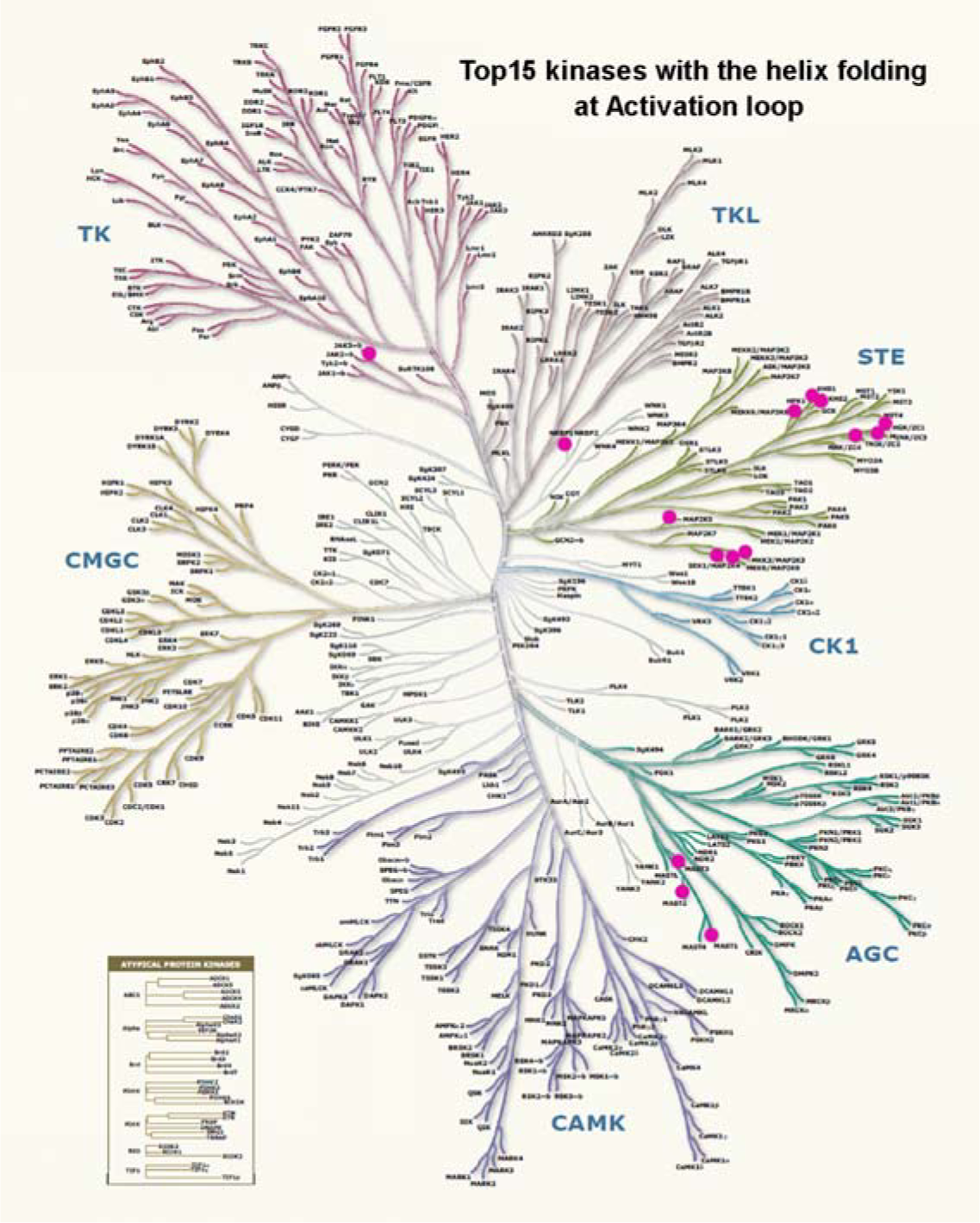

**Table 1.**
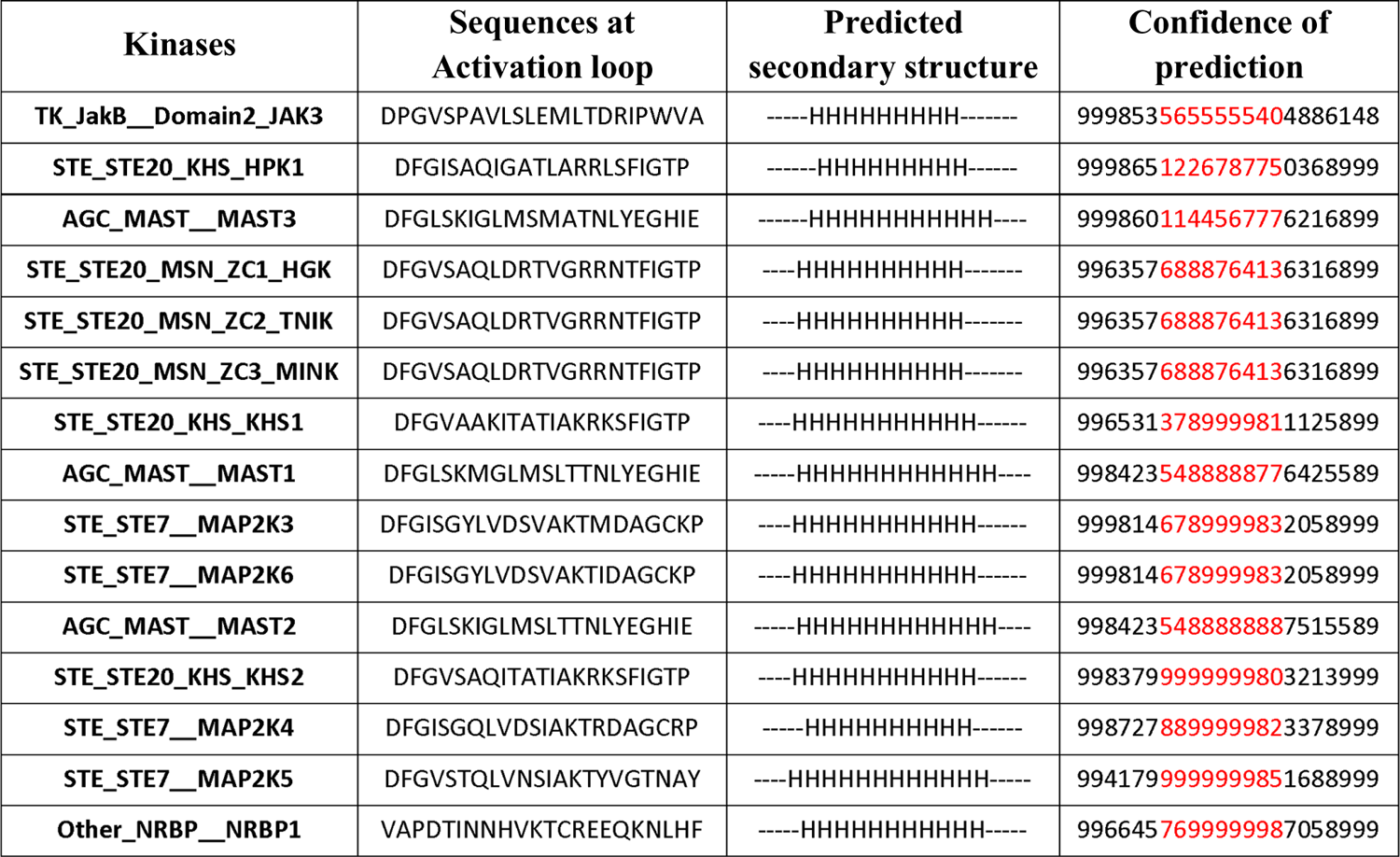
Top 15 kinases predicted with high confidence to have a helix at part of the activation loop (red color).

## Conclusions

Recently FDA approved type-III allosteric inhibitors provide an important opportunity to design and discover more efficient and lower-dose kinase inhibitors. Study of the characteristics of the type-III inhibitor-bound binding site provides structural insights into the design of new allosteric drugs. Here we study the characteristics of the MEK binding site, the chemical nature of the inhibitors that bind MEK, the characteristics of binding and the dynamic nature of the interaction. Further, based on all 3D kinase structures, we screened for potential allosteric inhibitor-bound binding sites which revealed that BRAF and P38a were similar to MEK. Finally, based on the distinctive helix character of the MEK activation loop [35] we predicted 15 kinases to potentially have the allosteric site needed to accommodate type-III allosteric inhibitors. In summary, our in silico analysis and prediction contributes to the understanding of the structure and ligand requirements for designing allosteric inhibitors.

## Methods

### Function-site interaction fingerprint scheme

Function-site Interaction Fingerprint (Fs-IFP) is a method to determine the functional site binding characteristics and to compare binding sites on a proteome scale [5]. Here we use Fs-IFP to reveal the binding characteristics of the MEK-inhibitor complex for all released MEK structures from the Protein Data Bank (PDB)[37]. In brief, firstly we downloaded all released 35 MEK structures from the PDB. Herein the 28 ligand-bound MEK structures formed the MEK structure dataset; then we aligned all the binding sites of these ligand-bound structures using SMAP [31–33] and encoded the Fs-IFP as published in an earlier paper [5] using the Pyplif software [38]. The end result for each structure is a representation of the interactions between every involved amino acid and the ligand using one-dimension bits.

### Pharmacophore modeling

Pharmacophore modeling was performed using Maestro in Schrodinger release 2016.02 [39]. Based on the MEK structure dataset, we collected the activity data of all ligands for MEK, the binding affinity data are shown in Table S1. Among them, 19 entries with MEK IC50 value were used as input. The pharmacophore model was trained using the ligands with an IC50 of less than 10 nM.

### MD simulation

Two MD simulations were performed with starting conformations taken from the PDB; pdb id 4an2 for the MEK-Cobimetinib complex, and the MEK apo structure from pdb id 3zls. Both initial conformations were prepared for MD simulations using the ACEMD protocol [40]. The protonation states of both systems were assigned as pH=7.0, similar to the cellular environment. Then every His state and every disulfide bond were checked to make sure they conformed to pH=7.0. The systems were solvated in a rectangular water box with at least a 12Å shell buffer away from any-solute atoms and charged ions were added to ensure an ionic strength of 0.20 M and electroneutrality. The CHARMM27 force field [41, 42], CHARMM general force field [43] and TIP3P force field were used for the kinase, ligand, and water molecules, respectively. Both systems were relaxed with a conversional MD protocol including 2ps minimization, 100ps for NVT, 1ns for NPT with heavy-atom constraints and 1ns for NPT without any constraints. Subsequently, 0.6 μs MD simulations were performed for every system. In both MD simulations all bonds were constrained using SHAKE and the integration time step was 4 fs. The temperature bath used the Langevin method, and 1atm pressure was maintained using the Berendsen method. Both simulations were carried out using the ACEMD software [40] on the NIH high-performance Biowulf cluster (<u>https://hpc.nih.gov/</u>). The MD results were analyzed using the conformations during the last 0.5 μs MD trajectory. The analysis of MD trajectories, including Root Mean Square Deviation (RMSD), Root Mean Square Fluctuation (RMSF) and Principal Component Analysis (PCA), were performed with the Wordom tool [44].

### Screening for similar binding pockets across the human structural kinome

Approximately three thousands protein kinase structures have been solved by X-ray and NMR methods. We assembled 2797 kinase structures from the PDB which included the catalytic domain as the human structural kinome [5]. We then used a MEK-Cobimetinib complex (PDB id 4lmn) as a template to rank similar binding sites from the human structural kinome by performing a one-to-all comparison using SMAP [31–33]. The top-ranked binding sites with *p*-values☐<☐0.05 were chosen for further analysis.

### Secondary structure prediction

Determining type-III inhibitors needs to be validated, not only by biological activity assay, but also through analysis of an appropriate crystal structure [8]. Unfortunately, the type-III binding pocket is often not present in the crystal structure when there is no specific type-III inhibitor co-crystalized [8]. To obtain the MEK-similar allosteric site in such kinases, we predicted the MEK-like secondary structure of the activation loop to establish the binding pose of the type-III inhibitor [35]. We used a protein secondary structure prediction server Jpred4 [45] for the task. First, the 516 kinase domain sequences for the eukaryotic protein kinase superfamily were downloaded at kinase.com and were aligned using the cluster omega software [46]. From the alignment the amino acid sequences of the activation loop for every kinase were extracted with additional 11 N-terminal and 11 C-terminal residues adjoining the DFG segments. Finally, the activation loop structure was predicted using the JPred RESTful API(v1.5) [45] with default parameters.

## Acknowledgments

This research was supported by the Intramural Research Program of the National Library of Medicine, National Institutes of Health (Z.Z. and P.E.B.), the National Library of Medicine, National Institutes of Health under Award R01LM011986 (L.X.), and the National Institute on Minority Health and Health Disparities, National Institutes of Health, under Award G12MD007599 (L.X.).

## Supporting information captions

**Fig S1.** ACP-Cobimetinib-MEK co-crystal structure (pdb id 4an2). a) The cartoon modeling. b) 2D diagrams for Cobimetinib/MEK interactions including ACP (marked as Acp1383) by LigPlot+.

**Fig S2.** The Ca-atom RMSD.

**Fig S3.** Sequence level similarity for the three kinases (BRAF, MEK, and P38a).

**Table S1.** List of active inhibitors for MEK.

## References

1. Klebl B, Müller G, Hamacher M, editors. Protein Kinase as Drug Targets. Weinheim: WILEY-VCH Verlag GambH & Co. KGaA; 2011.

2. Manning G, Whyte DB, Martinez R, Hunter T, Sudarsanam S. The protein kinase complement of the human genome. Science. 2002;298(5600):1912–34. doi: 10.1126/science.1075762. PubMed PMID: 12471243.

3. Lahiry P, Torkamani A, Schork NJ, Hegele RA. Kinase mutations in human disease: interpreting genotype-phenotype relationships. Nat Rev Genet. 2010;11(1):60–74. doi: 10.1038/nrg2707. PubMed PMID: 20019687.

4. Abramson R. Overview of Targeted Therapies for Cancer. My Cancer Genome <u>https://www.mycancergenome.org/content/molecular-medicine/overview-of-targeted-therapies-for-cancer/</u> (Updated April 26). 2016.

5. Zhao Z, Xie L, Xie L, Bourne PE. Delineation of Polypharmacology across the Human Structural Kinome Using a Functional Site Interaction Fingerprint Approach. J Med Chem. 2016;59(9):4326–41. doi: 10.1021/acs.jmedchem.5b02041. PubMed PMID: 26929980; PubMed Central PMCID: PMC4865454.

6. Wu P, Nielsen TE, Clausen MH. Small-molecule kinase inhibitors: an analysis of FDA-approved drugs. Drug Discov Today. 2016;21(1):5–10. doi: 10.1016/j.drudis.2015.07.008. PubMed PMID: 26210956.

7. Rask-Andersen M, Zhang J, Fabbro D, Schioth HB. Advances in kinase targeting: current clinical use and clinical trials. Trends Pharmacol Sci. 2014;35(11):604–20. doi: 10.1016/j.tips.2014.09.007. PubMed PMID: 25312588.

8. Muller S, Chaikuad A, Gray NS, Knapp S. The ins and outs of selective kinase inhibitor development. Nat Chem Biol. 2015;11(11):818–21. doi: 10.1038/nchembio.1938. PubMed PMID: 26485069.

9. Gharwan H, Groninger H. Kinase inhibitors and monoclonal antibodies in oncology: clinical implications. Nat Rev Clin Oncol. 2016;13(4):209–27. doi: 10.1038/nrclinonc.2015.213. PubMed PMID: 26718105.

10. Kooistra AJ, Kanev GK, van Linden OP, Leurs R, de Esch IJ, de Graaf C. KLIFS: a structural kinase-ligand interaction database. Nucleic Acids Res. 2016;44(D1):D365–71. doi: 10.1093/nar/gkv1082. PubMed PMID: 26496949; PubMed Central PMCID: PMC4702798.

11. Roskoski R, Jr. Classification of small molecule protein kinase inhibitors based upon the structures of their drug-enzyme complexes. Pharmacol Res. 2016;103:26–48. doi: 10.1016/j.phrs.2015.10.021. PubMed PMID: 26529477.

12. Gushwa NN, Kang S, Chen J, Taunton J. Selective targeting of distinct active site nucleophiles by irreversible SRC-family kinase inhibitors. J Am Chem Soc. 2012;134(50):20214–7. doi: 10.1021/ja310659j. PubMed PMID: 23190395; PubMed Central PMCID: PMC3729023.

13. Zhao Z, Wu H, Wang L, Liu Y, Knapp S, Liu Q, et al. Exploration of type II binding mode: A privileged approach for kinase inhibitor focused drug discovery? ACS Chem Biol. 2014;9(6):1230–41. doi: 10.1021/cb500129t. PubMed PMID: 24730530; PubMed Central PMCID: PMC4068218.

14. Rice KD, Aay N, Anand NK, Blazey CM, Bowles OJ, Bussenius J, et al. Novel Carboxamide-Based Allosteric MEK Inhibitors: Discovery and Optimization Efforts toward XL518 (GDC-0973). ACS Med Chem Lett. 2012;3(5):416–21. doi: 10.1021/ml300049d. PubMed PMID: 24900486; PubMed Central PMCID: PMC4025802.

15. Hatzivassiliou G, Haling JR, Chen H, Song K, Price S, Heald R, et al. Mechanism of MEK inhibition determines efficacy in mutant KRAS-versus BRAF-driven cancers. Nature. 2013;501(7466):232–6. doi: 10.1038/nature12441. PubMed PMID: 23934108.

16. Wu P, Clausen MH, Nielsen TE. Allosteric small-molecule kinase inhibitors. Pharmacol Ther. 2015;156:59–68. doi: 10.1016/j.pharmthera.2015.10.002. PubMed PMID: 26478442.

17. Hindie V, Stroba A, Zhang H, Lopez-Garcia LA, Idrissova L, Zeuzem S, et al. Structure and allosteric effects of low-molecular-weight activators on the protein kinase PDK1. Nat Chem Biol. 2009;5(10):758–64. doi: 10.1038/nchembio.208. PubMed PMID: 19718043.

18. Yap TA, Yan L, Patnaik A, Fearen I, Olmos D, Papadopoulos K, et al. First-in-man clinical trial of the oral pan-AKT inhibitor MK-2206 in patients with advanced solid tumors. J Clin Oncol. 2011;29(35):4688–95. doi: 10.1200/JCO.2011.35.5263. PubMed PMID: 22025163.

19. Jia Y, Yun CH, Park E, Ercan D, Manuia M, Juarez J, et al. Overcoming EGFR(T790M) and EGFR(C797S) resistance with mutant-selective allosteric inhibitors. Nature. 2016;534(7605):129–32. doi: 10.1038/nature17960. PubMed PMID: 27251290; PubMed Central PMCID: PMC4929832.

20. Bourne PE, Xie L. Harnessing ‘Big Data’ in Systems Pharmacology. Annu Rev Pharmacol Toxicol. 2017;57(1):doi:10.1146/annurev-pharmtox-010716-104659. doi: doi:10.1146/annurev-pharmtox-010716-104659.

21. Zhao Z, Martin C, Fan R, Bourne PE, Xie L. Drug repurposing to target Ebola virus replication and virulence using structural systems pharmacology. BMC Bioinformatics. 2016;17:90. doi: 10.1186/s12859-016-0941-9. PubMed PMID: 26887654; PubMed Central PMCID: PMC4757998.

22. De Vivo M, Masetti M, Bottegoni G, Cavalli A. Role of Molecular Dynamics and Related Methods in Drug Discovery. J Med Chem. 2016;59(9):4035–61. doi: 10.1021/acs.jmedchem.5b01684. PubMed PMID: 26807648.

23. Garnock-Jones KP. Cobimetinib: First Global Approval. Drugs. 2015;75(15):1823–30. doi: 10.1007/s40265-015-0477-8. PubMed PMID: 26452567.

24. Smith CA, Rayment I. Active site comparisons highlight structural similarities between myosin and other P-loop proteins. Biophys J. 1996;70(4):1590–602. doi: 10.1016/s0006-3495(96)79745-x.

25. Deyrup AT. Deletion and Site-directed Mutagenesis of the ATP-binding Motif (P-loop) in the Bifunctional Murine Atp-Sulfurylase/Adenosine 5'-Phosphosulfate Kinase Enzyme. J Biol Chem. 1998;273(16):9450–6. doi: 10.1074/jbc.273.16.9450.

26. McClendon CL, Kornev AP, Gilson MK, Taylor SS. Dynamic architecture of a protein kinase. Proc Natl Acad Sci USA. 2014;111(43):E4623–31. doi: 10.1073/pnas.1418402111. PubMed PMID: 25319261; PubMed Central PMCID: PMC4217441.

27. Mazanetz MP, Laughton CA, Fischer PM. Investigation of the flexibility of protein kinases implicated in the pathology of Alzheimer's disease. Molecules. 2014;19(7):9134–59. doi: 10.3390/molecules19079134. PubMed PMID: 24983862.

28. Adams JA. Kinetic and Catalytic Mechanisms of Protein Kinases. Chem Rev. 2001;101(8):2271–90. doi: 10.1021/cr000230w.

29. Roskoski R, Jr. MEK1/2 dual-specificity protein kinases: structure and regulation. Biochem Biophys Res Commun. 2012;417(1):5–10. doi: 10.1016/j.bbrc.2011.11.145. PubMed PMID: 22177953.

30. Kornev AP, Haste NM, Taylor SS, Eyck LF. Surface comparison of active and inactive protein kinases identifies a conserved activation mechanism. Proc Natl Acad Sci USA. 2006;103(47):17783–8. doi: 10.1073/pnas.0607656103. PubMed PMID: 17095602; PubMed Central PMCID: PMC1693824.

31. Xie L, Bourne PE. A robust and efficient algorithm for the shape description of protein structures and its application in predicting ligand binding sites. BMC Bioinformatics. 2007;8 Suppl 4:S9. doi: 10.1186/1471-2105-8-S4-S9. PubMed PMID: 17570152; PubMed Central PMCID: PMC1892088.

32. Xie L, Bourne PE. Detecting evolutionary relationships across existing fold space, using sequence order-independent profile-profile alignments. Proc Natl Acad Sci USA. 2008;105(14):5441–6. doi: 10.1073/pnas.0704422105. PubMed PMID: 18385384; PubMed Central PMCID: PMC2291117.

33. Xie L, Xie L, Bourne PE. A unified statistical model to support local sequence order independent similarity searching for ligand-binding sites and its application to genome-based drug discovery. Bioinformatics. 2009;25(12):i305–12. doi: 10.1093/bioinformatics/btp220. PubMed PMID: 19478004; PubMed Central PMCID: PMC2687974.

34. Adachi-Yamada T, Nakamura M, Irie K, Tomoyasu Y, Sano Y, Mori E, et al. p38 mitogen-activated protein kinase can be involved in transforming growth factor beta superfamily signal transduction in Drosophila wing morphogenesis. Mol Cell Biol. 1999;19(3):2322–9. PubMed PMID: 10022918; PubMed Central PMCID: PMC84024.

35. Lee CC, Jia Y, Li N, Sun X, Ng K, Ambing E, et al. Crystal structure of the ALK (anaplastic lymphoma kinase) catalytic domain. Biochem J. 2010;430(3):425–37. doi: 10.1042/BJ20100609. PubMed PMID: 20632993.

36. Matsumoto T, Kinoshita T, Kirii Y, Yokota K, Hamada K, Tada T. Crystal structures of MKK4 kinase domain reveal that substrate peptide binds to an allosteric site and induces an auto-inhibition state. Biochem Biophys Res Commun. 2010;400(3):369–73. doi: 10.1016/j.bbrc.2010.08.071. PubMed PMID: 20732303.

37. Berman HM, Bhat TN, Bourne PE, Feng Z, Gilliland G, Weissig H, et al. The Protein Data Bank and the challenge of structural genomics. Nat Struct Biol. 2000;7 Suppl:957–9. doi: 10.1038/80734. PubMed PMID: 11103999.

38. Radifar M, Yuniarti N, Istyastono EP. PyPLIF: python-based protein-ligand fnteraction fingerprinting. Bioinformatics. 2013;9(6):325–8. doi: 10.6026/97320630009325. PubMed PMID: 23559752; PubMed Central PMCID: PMC3607193.

39. Schrödinger Release 2016-2: Maestro, version 10.6, Schrödinger, LLC, New York, NY, 2016.

40. Harvey MJ, Giupponi G, Fabritiis GD. ACEMD: Accelerating Biomolecular Dynamics in the Microsecond Time Scale. J Chem Theory Comput. 2009;5(6):1632–9. doi: 10.1021/ct9000685. PubMed PMID: 26609855.

41. Brooks BR, Bruccoleri RE, Olafson BD, States DJ, Swaminathan S, Karplus M. CHARMM: A program for macromolecular energy, minimization, and dynamics calculations. J Comput Chem. 1983;4(2):187–217. doi: 10.1002/jcc.540040211.

42. Brooks BR, Brooks CL, 3rd, Mackerell AD, Jr., Nilsson L, Petrella RJ, Roux B, et al. CHARMM: the biomolecular simulation program. J Comput Chem. 2009;30(10):1545–614. doi: 10.1002/jcc.21287. PubMed PMID: 19444816; PubMed Central PMCID: PMC2810661.

43. Vanommeslaeghe K, Hatcher E, Acharya C, Kundu S, Zhong S, Shim J, et al. CHARMM general force field: A force field for drug-like molecules compatible with the CHARMM all-atom additive biological force fields. J Comput Chem. 2010;31(4):671–90. doi: 10.1002/jcc.21367. PubMed PMID: 19575467; PubMed Central PMCID: PMC2888302.

44. Seeber M, Cecchini M, Rao F, Settanni G, Caflisch A. Wordom: a program for efficient analysis of molecular dynamics simulations. Bioinformatics. 2007;23(19):2625–7. doi: 10.1093/bioinformatics/btm378. PubMed PMID: 17717034.

45. Drozdetskiy A, Cole C, Procter J, Barton GJ. JPred4: a protein secondary structure prediction server. Nucleic Acids Res. 2015;43(W1):W389–94. doi: 10.1093/nar/gkv332. PubMed PMID: 25883141; PubMed Central PMCID: PMC4489285.

46. Li W, Cowley A, Uludag M, Gur T, McWilliam H, Squizzato S, et al. The EMBL-EBI bioinformatics web and programmatic tools framework. Nucleic Acids Res. 2015;43(W1):W580–4. doi: 10.1093/nar/gkv279. PubMed PMID: 25845596; PubMed Central PMCID: PMC4489272.

